# Frontal theta synchronization facilitates the updating of statistical regularities, evidenced by predictive eye movements

**DOI:** 10.1101/2025.08.18.670954

**Authors:** Adrienn Holczer, Orsolya Pesthy, Teodóra Vékony, Gábor Csifcsák, Dezső Németh

## Abstract

Frontal midline theta oscillations are key neural markers for learning, set-shifting, and adaptive behavior, signaling cognitive control and the reorganization of neural representations. The present study explored how these oscillations mediate the extraction and updating of statistical regularities. We delivered 6 Hz in-phase transcranial alternating current stimulation (tACS) or sham tACS, synchronizing frontal midline theta during an eye-tracking probabilistic sequence learning task designed to test cognitive flexibility and assess pre-stimulus gaze direction changes. A novel probabilistic sequence with a partially overlapping structure was introduced that allowed us to distinguish between the retention of old sequences and the acquisition of new ones. Following comparable statistical learning in both groups during the stimulation session, our results showed that tACS reduced the erroneous anticipations of previously learned regularities, and allowed participants to show anticipations corresponding to the previously learnt regularities while being able to anticipate novel regularities flexibly. These results suggest a role of frontal midline theta in the flexible rewiring of the mental representations of prior probabilistic structures, and in making predictions more accurate.

## Introduction

Adaptation to a constantly changing environment requires the effective updating of neural representations through neuroplasticity (Zhong et al. 2025). While most neuroplastic changes have long been attributed to task-specific training, recent findings highlight the significance of unsupervised learning, emphasizing that unrewarded exposure to stimuli can reshape internal models and facilitate a subsequent learning process (Zhong et al. 2025). The ability to implicitly extract predictable regularities by mere exposure from a noisy environment, known as statistical learning, is fundamental to building and updating our mental models (Turk-Browne et al. 2010; Aslin 2017; Conway 2020), and is involved in acquiring various cognitive, language, social, and motor skills (Daw et al. 2005; Horváth et al. 2022). Statistical learning relies primarily on model-free, stimulus-driven processes, and evidence suggests that strong engagement of model-based, top-down mechanisms such as cognitive control or working memory can even hinder it (Virag et al. 2015; Janacsek and Nemeth 2022; Pedraza et al. 2024; Pedraza et al. 2025; Pesthy et al. 2025). More specifically, frontal midline theta oscillations are considered a key neural signature for learning, set-shifting, and adaptive behavior, signaling the need for cognitive control and the reorganization of neural representations (Enriquez-Geppert et al. 2014; Eisma et al. 2021; Verbeke et al. 2021). Thus, investigating the neural mechanisms underlying statistical learning—particularly the role of frontal midline theta oscillations in mediating the extraction, and most importantly, updating of statistical regularities—is essential.

Administering transcranial alternating current stimulation (tACS) is a widely used tool to investigate causal relationships between theta oscillations and various cognitive processes (Antal and Paulus 2013). Theta tACS has been shown to improve cognitive control, working memory, and cognitive flexibility; moreover, it resulted in an increase of related neuropsychological correlates (Fusco et al. 2018; Rostami et al. 2021; Zhu et al. 2023; Debnath et al. 2025). Despite these promising effects in related domains, the literature examining the role of theta tACS in statistical learning is limited. So far, only a few studies have investigated its impact on statistical learning specifically, and these typically reported no performance change (Zavecz et al. 2020; Diedrich et al. 2024). However, frontal midline theta activity during statistical learning has been suggested to regulate the involvement of top-down processing (Lum et al. 2024). This highlights the need for further investigation into how theta tACS might influence the robustness or flexibility of statistical learning processes. The present study implements some methodological advances, including the use of eye-tracking to assess predictive processing and the re-assessment of statistical learning following a delay.

To dissociate momentary performance and the underlying knowledge of statistical regularities, it is insufficient to measure performance only during active practice (Kiss et al. 2022). Learning is a dynamic process that unfolds over time, relying on offline periods to stabilize memory traces (Janacsek and Nemeth 2012; Fanuel et al. 2022). Furthermore, a robust test of learning is its resilience in a changing environment, requiring the maintenance of prior knowledge in the face of interference. Additionally, flexibly updating prior knowledge with new information demonstrates the effectiveness of adaptation. Considering these core features of learning, we identified two critical dimensions that previous studies on the relationship of frontal midline theta and statistical learning have failed to address. Specifically, they have neglected: (1) re-testing statistical learning following a delay to capture tACS aftereffects and the stability of memory traces, and (2) assessing the updating of acquired knowledge against new, interfering information.

Supporting the need for delayed testing following the active practice during tACS, a meta-analysis has suggested that tACS effects may manifest more prominently offline (i.e., when performance is measured following the stimulation) (Grover et al. 2023). Another related study, which involved disrupting prefrontal cortex function by transcranial magnetic stimulation, found changes in statistical learning only after a delay, further supporting the idea that the mechanism of action may involve offline neuroplastic processes (Janacsek et al. 2015; Ambrus et al. 2020). To address this, the present study introduces a post-tACS session to specifically assess the delayed effects of the stimulation. This step would overcome the limitation of Zavecz et al. (2020), where only the online effects of tACS (i.e., the performance during the stimulation) were assessed. In another study, Diedrich et al. (2024) assessed the effect of tACS on the performance on a probabilistic sequence learning task following a delay; however, none of the studies mentioned above have directly tested whether theta tACS can alter the acquisition of an interfering statistical regularity after gaining stable knowledge of an original sequence (Zavecz et al. 2020; Diedrich et al. 2024; Lum et al. 2024). By adding interfering information following the initial learning phase, we aim to investigate the resistance of the acquired knowledge as well as the effectiveness of updating and acquiring a novel but similar probabilistic structure.

In the present study, to address the role of frontal midline theta in statistical learning, especially in the updating of already acquired statistical regularities with novel information, we combine 6 Hz tACS synchronizing frontal midline theta with a probabilistic sequence learning paradigm designed to test cognitive flexibility. We introduce a novel probabilistic sequence containing a partially new structure, allowing us to dissociate the retention of an old sequence from the acquisition of a new one. Critically, we move beyond the limitations of simple reaction times by using eye-tracking to implement a highly sensitive method that assesses the mechanisms of predictive processing through examining pre-stimulus gaze directions (Zolnai et al. 2022), providing insight into how learned statistical regularities are deployed and updated.

## Methods

### Participants

We recruited healthy young adults with normal or corrected-to-normal vision. None of the participants reported a history of any neurological or psychiatric disorders or the use of any drugs affecting the function of the central nervous system. Participants had to meet the safety restrictions of tACS (no history of epilepsy, having a first-degree relative with epilepsy, previous head injury or concussion, metallic implants in the cephalic region, or any implanted electronic devices like pacemakers). Participants provided written informed consent before enrollment. The experimental protocol was approved by the local Ethics Committee (2018/52). The experiment was conducted in accordance with the Declaration of Helsinki, as revised in 2008.

One hundred and seven participants were recruited for the study. Participants were assigned to one of the following three groups using computer-generated random allocation: synchronization (33 participants), desynchronization (35 participants), or sham stimulation (39 participants, with the electrode montage identical to that of the desynchronization group). Participants were not aware of their group assignment.

Due to an irreparable failure of the eye-tracking device, data collection was halted. This failure did not affect the quality of the previously collected data. Due to the unequal number of participants excluded from the analysis because of corrupted/missing data or experimenter error (see Supplementary Methods), and since further data collection was not possible, the data from the desynchronization group were not analyzed since the insufficient group size prevented the generation of reliable estimates. We further excluded the data of six participants (one from the sham group and five from the synchronization group) because they did not meet our eye-tracking data quality requirements (see Supplementary Methods). Overall, the final sample comprised data from 53 participants (38 female, *M*_age_ = 21.42 years, *SD*_age_ = 2.61 years), assigned to either the synchronization group (final group size: 21) or the sham group (final group size: 32 participants).

To maximize statistical power, all participants with valid data were included in the analysis of the resting-state electroencephalography (EEG) recordings and the control episodic memory task when we compared the results per session, independently of the main probabilistic sequence learning task. Overall, the data of 34 participants from the sham group and 26 from the synchronization group were included for the episodic memory task. For the resting-state EEG, the data of 31 participants from the sham group and 29 participants from the synchronization group were used after removing the data with experimenter error or corrupted data.

### Alternating serial reaction time (ASRT) task

Statistical learning was assessed using a modified version of the Alternating Serial Reaction Time (ASRT) task using eye-tracking (Zolnai et al. 2022). A Tobii Pro X3-120 eye tracker was used with a 120 Hz sampling rate (Tobii 2017). The device was mounted to the bottom of a 24-inch AOC LED monitor with a resolution of 1920 × 1080. Data acquisition was carried out using the Tobii Pro Python SDK (Tobii Pro SDK for Python 2020), integrated with a PsychoPy-based experimental script (version: 3.2.3, Peirce et al. 2019). An approximate subject-screen distance of 65 cm was maintained throughout the task. Gaze position estimation and fixation identification were performed as described in (Zolnai et al. 2022).

During the ASRT task, four empty circles were presented to the participants—one in each corner of a 1,920 × 1,080 resolution screen. The diameter of the circles was 3 cm (≈ 2.64° visual angle). The four stimulus positions were spaced equally apart (15 cm, ≈ 13.16° visual angle) and from the center of the screen (10.6 cm, ≈ 9.32° visual angle).

A trial consisted of the presentation of a target stimulus, defined as a single circle turning blue (Figure 1, Panel A). Participants were instructed to look at the target stimulus as quickly as possible. Once participants fixated on the target for at least 100 ms, it was considered a response and the stimulus disappeared. The next target stimulus was presented with a response–stimulus interval of 500 ms. Trials were organized into blocks of 85 trials. Between blocks, there was a pause where participants received feedback on their speed. Solely for data analysis purposes, each five blocks was further nested into epochs. Note that the breaks did not differ between blocks and epochs. Before the actual ASRT task, all participants underwent one practice epoch (i.e., five blocks, each with 85 trials) of randomly appearing trials at the beginning of the experiment to familiarize themselves with the task.

**Figure 1.**
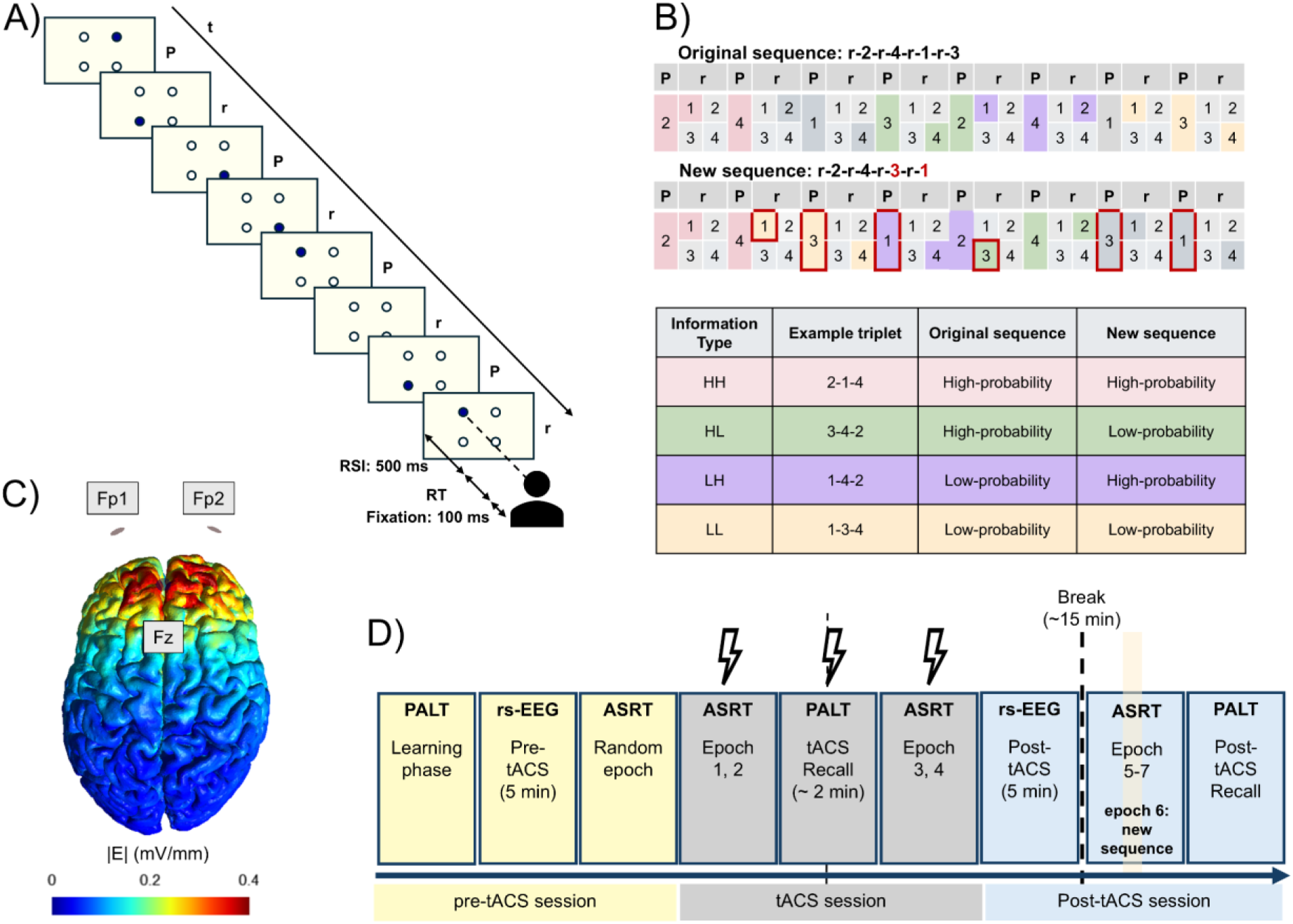
Task design and experimental paradigm. (A) The target stimulus marked in dark blue could appear in one of the four possible locations. Participants were instructed to focus on the target stimulus until it disappeared. Pattern (P) and random (r) stimuli alternated. (B) Examples for the original and the new sequence. There was a partial overlap between the two sequences (marked by the original color), while the elements marked by red changed in the new sequence. This partial overlapping structure resulted in some triplets remaining high-probability according to both sequences (HH, marked with pink), some that were high-probability became low-probability in the new sequence (HL, marked with green), some previously low-probability triplets turned high-probability in the new sequence (LH, marked with purple), and some remained low-probability according to both sequences (LL, marked with yellow). (C) Simulation of the electric currents in the Synchronization group using the SimNIBS software. 6 Hz in-phase tACS was delivered to electrodes over the Fp1 and Fp2 with a return electrode over Fz. (D) Study design. Participants first completed the learning phase of the Paired Associate Learning Task (PALT). Following a 5-minute resting-state EEG, participants completed a random epoch that consisted of randomized trials. During the tACS, participants completed in order: epoch 1-2 of the ASRT task, a retrieval session of the PALT, and epoch 3-4 of the ASRT task. A second resting-state EEG was followed by a 15-minute break, after which participants were reintroduced to the original sequence in epoch 5, to a previously unseen new sequence in epoch 6, and finally, in epoch 7, the original sequence was presented again. *Note. P: pattern, r: random, RSI: response-to-stimulus interval, RT: reaction time, HH: high-high, HL: high-low, LH: low-high, LL: low-low, PALT: Paired Associate Learning Task, rs-EEG: resting-state electroencephalography, ASRT: alternating serial reaction time task*

Unbeknownst to the participants, stimuli appeared in a predefined probabilistic eight-element sequence in which pattern and random elements were alternating (e.g., r-1-r-2-r-3-r-4, where numbers indicate the location of one of the four placeholders on the screen (1 = lower left, 2 = lower right, 3 = upper right, 4 = upper left), and “r” denotes a random location). Because of the structure of alternating elements, some runs of three consecutive trials (i.e., triplets) were more predictable (high-probability triplets) than others (low-probability triplets). Importantly, each trial (i.e., the presentation of one target stimulus) can be considered the third element of a high- or low-probability triplet based on its predictability by the trial n-2. This is because the element n of a given high-probability triplet is more predictable based on element n-2 of the triplet due to second-order transitional probabilities, whereas, in low-probability triplets, this predictability is weaker.

Considering r-2-r-4-r-1-r-3 as an example, high-probability triplets can either be formed by two pattern elements and a random item in between (comprising 50% of all trials) or by two random elements and a pattern element if it fits the ASRT sequence (comprising 12.5% of all trials), e.g., 2-X-**4**, 4-X-**1**, where X indicates the middle element of the triplet, regardless of whether it is a random element or part of the pattern. Low-probability triplets are those triplets that cannot be formed by the pattern (e.g., 2-X-**1**, 4-X-**3**, comprising the remaining 37.5% of all trials). As the task progresses, participants tend to respond faster to trials that are more probable (i.e., last items of a high-probability triplet) than those that are less likely to appear (i.e., last item of a low-probability triplet). Probabilistic sequence learning in terms of reaction time is the difference between the last items of high- and low-probability triplets (i.e., high- or low-probability trials). A greater difference between high- and low-probability trials indicates better probabilistic sequence learning.

### Pre-stimulus gaze direction monitoring

Using the eye-tracking version of the ASRT task, it was also possible to record eye movements during the response-stimulus interval. We followed the protocol of (Zolnai et al. 2022) in analyzing eye movements. We divided the screen into four equal imaginary fields by the center lines. Each of these four fields contained one of the possible placeholders for the target stimulus. We used the last valid gaze position of the response-to-stimulus interval, i.e., before the new stimulus appeared, to identify whether the participant moved their gaze to one of the three other fields from the one where the last target stimulus was presented. If so, we considered that a *pre-stimulus gaze direction change*. If the final registered pre-stimulus gaze direction corresponded to the last element of a high-probability triplet rather than to low-probability triplets, we marked it as a *learning-dependent anticipatory eye movement,* indicating that participants learned the probabilistic sequence of the ASRT task (e.g., in case of the 2-1-4 high-probability triplet, if after 1 disappeared, the last valid gaze position of the participant was recorded at the quadrant where 4 would be presented, it was considered a learning-dependent anticipatory eye movement). Notably, due to the statistical nature of the task, learning-dependent anticipations do not necessarily mean that the prediction is correct and the target stimulus appears over that placeholder. Furthermore, we calculated the *learning-dependent anticipation ratio* by dividing the amount of learning-dependent anticipatory eye movements by the amount of pre-stimulus gaze direction changes.

Overall, participants showed a pre-stimulus gaze direction change in 21.67% of all trials in our task. In 6.80% of all trials, the pre-stimulus gaze direction corresponded to a high-probability triplet; thus, they were considered ***learning-dependent anticipations***. The ***learning-dependent anticipation ratio,*** i.e., the ratio of learning-dependent and not-learning-dependent anticipations, was 31.38%.

We also decided to employ a more nuanced analysis of pre-stimulus gaze directions by categorizing each trial into four possible ***pre-stimulus gaze direction types***. We created four categories, distinguishing whether the participant anticipated a high or low-probability triplet, and whether that anticipation matched the actual outcome. All pre-stimulus gaze direction changes were organized into one of the following categories: 1) *Learning-Dependent Correct* (LDC) refers to the anticipation and following appearance of a high-probability triplet, 2) *Learning-Dependent Error* (LDE) is the anticipation of a high-probability triplet, when a low-probability triplet appears, 3) *Not-Learning-Dependent Correct* (NLDC) is defined as a pre-stimulus gaze direction change toward a quadrant of the screen where a low-probability triplet may appear when a low-probability triplet actually appears, and 4) *Not-Learning-Dependent Error* (NLDE) when the pre-stimulus gaze direction is at a low-probability triplet’s position, but the target stimulus appears at another location.

Rather than simply accounting for learning-dependent or not-learning-dependent anticipations, we integrated both the probability structure of the task, by weighing each type of anticipatory eye movement according to its chance-level probability (LDC: 0.15625, LDE: 0.09375, NLDC: 0.09375, NLDE: 0.65625, see Supplementary Methods, Supplementary Table S1), and the actual accuracy of the anticipations. This approach allows us to more sensitively capture learning-related changes in pre-stimulus gaze directions, taking into account both the likelihood of a given anticipation and whether it matched the upcoming stimulus. Moreover, it may provide insight into error-related responses as well as the sensitivity of participants regarding the statistical probabilities of specific types of stimuli, while accounting for the baseline probability of each type.

### Introducing a new sequence in the ASRT task

To assess resistance to interference and the acquisition of a new probabilistic structure, we introduced a new, partially overlapping sequence during the post-tACS phase of the experiment (Figure 1, Panel B). Considering r-2-r-4-r-1-r-3 as the original sequence, an example of a partially overlapping sequence would be r-2-r-4-r-3-r-1. In the new sequence, the first and the second pattern elements remain at the original locations, while the positions of the third and fourth pattern elements have changed. Due to this change in the sequence, each stimulus could be labeled as one of the following: *HH* (“*high-high*” triplets) if it was the last element of an originally high-probability triplet that remained high probability according to the new sequence; *HL* (“*high-low*”) if it was the last element of a previously high-probability triplet that turned into low-probability triplet in the interference blocks, i.e., could be considered as a measure of old knowledge acquired during the tACS session; *LH* (“*low-high*” triplets), when the originally low-probability triplet turned into high-probability triplet, thus, can be considered as the acquisition of new information; and *LL* (“*low-low*” triplets) if they remained low probability according to both sequences.

Four of the originally high-probability triplets remained high probability (HH). Twelve previously high-probability triplets according to the original sequence turn into low-probability triplets (HL). Of the 48 originally low-probability triplets, 12 turn into high-probability triplets (LH), and 36 remain low probability in both sequences (LL).

### Paired Associate Learning Task (PALT)

The PALT served as a memory control task to assess declarative/episodic learning. The PALT was presented using the E-Prime 3.0 software (Psychology Software Tools, Pittsburgh, PA). The task consisted of a learning phase and two phases of recollection. During the learning phase, participants were presented with 23 pairs of schematic drawings of animals and objects. Each pair consisted of an image of an animal and an object presented side by side on the computer screen. Participants were instructed to verbally name both images. Participants were not aware that this was a memory task.

During the retrieval phases, participants were shown a total of 32 pairs (24 in the tACS retrieval and 8 in the post-tACS retrieval) and instructed to indicate verbally 1) whether each individual image was presented during the learning phase and 2) whether the pair was the same as presented in the learning phase. The structure of the retrieval phase was the following: image pairs were either 1) the image pair was the same as in the learning phase (old-old, original; 8 pairs), 2) both images were presented in the learning phase but the pairing was different (old-old, rearranged; 8 pairs), 3) only one of the images was presented during the learning phase, the other was a novel image (old-new; 8 pairs), or 4) both images were novel and not presented in the learning phase (new-new; 8 pairs). The experimenter coded the participants’ verbal responses and operated the program by pressing the corresponding key on the keyboard. A 500-ms fixation cross was displayed between each stimulus presentation.

Three indices are calculated based on the responses: 1) ***associate learning score***, considered the primary outcome measure of the task, referring to whether participants were able to differentiate original old-old image pairs from rearranged old-old pairs, 2) ***item memory score***, which indicates if participants were able to differentiate already seen image pairs from completely new pairs, and 3) ***recollection score***, indicating if participants were able to differentiate pairs they saw together from pairs they saw separately. We calculated the associate learning score by subtracting the ratio of correct responses to rearranged old-old responses (hit rate) from the ratio of correct responses to old-old original responses (hit rate). The item memory score was calculated by subtracting the ratio of incorrect old-old responses to new-new pairs (false alarm) from the ratio of correct responses to old-old rearranged pairs (hit rate). The recollection score was calculated by subtracting the ratio of incorrect old-old original responses to old-old rearranged pairs (false alarm) from the ratio of correct responses to old-old original pairs (hit rate).

### Transcranial alternating current stimulation

The Startstim-20 device was used to deliver 6 Hz 2 mA peak-to-baseline stimulation (Neuroelectrics; www.neuroelectrics.com). The electrode montage and current intensity were selected based on preliminary simulations using the SimNIBS software (version 3.2.3, Thielscher et al. 2015) with the goal of reaching a current density of at least 0.4 V/m under the target region (Filmer et al. 2020). The active stimulation was administered for 20 minutes, while the sham stimulation lasted for 20 seconds. In the synchronization group, in-phase stimulation was delivered between the two stimulation electrodes positioned over Fp1 and Fp2, and a reference electrode was placed over Fz, according to the 10-20 EEG system (see Figure 1, Panel C). The electrode montage used for the desynchronization group was employed to deliver the sham stimulation (anti-phase stimulation was delivered between frontal stimulation electrodes: Fp1, Fp2, return electrode: FCz, and parietal stimulation electrodes: P3, P4, return electrode: CPz). We chose this montage to maximize scalp sensations as the blinding effectiveness of sham protocols has been questioned several times in the past (Sheffield et al. 2022). Such a brief electrical stimulation has not been linked to any relevant changes in cortical excitability (Dissanayaka et al. 2017) or EEG power (Pahor and Jaušovec 2018; Hosseinian et al. 2021).

### EEG recording and preprocessing

The Startstim-20 device was used to record resting-state EEG before and after the stimulation. Twenty electrodes were positioned according to the international 10-20 EEG system. Participants sat comfortably and were kept awake during the recordings. The recording lasted 5 minutes (the first 2.5 minutes in an eyes-open condition and 2.5 minutes with eyes closed). During the eyes-open condition, participants were asked to focus on a fixation cross in the middle of the computer screen. During acquisition, electrodes were referenced to a reference on the right earlobe. All electrode contact impedances were kept below 10 kΩ. EEG data were recorded with a sampling rate of 500 Hz.

EEG preprocessing was performed in Python (version 3.13.3) using the MNE package (version 1.9.0; Gramfort 2013). Recordings were band-pass filtered (0.5–40 Hz) with a FIR filter and a 50 Hz notch filter. Channels exhibiting persistent artifacts or low signal quality were interpolated using spherical splines after visual inspection. Next, independent component analysis (ICA) was performed to identify eye movement and other muscular artifacts using the MNE package. Independent components representing eye movements, up to a maximum of two components, were semi-automatically detected and identified by inspecting waveforms and topographical distributions. Subsequently, the data were re-referenced to the average of all channels. The continuous recording was segmented into 1-second-long segments, and baseline correction for the whole segment was used to remove DC offsets. Finally, segments with an amplitude deviation exceeding ±150 μV were rejected. Power spectral density was estimated by Fast Fourier transformation using Welch’s method with 1-second windows (yielding 1 Hz frequency resolution) separately for eyes open and eyes closed recordings. For each participant, power spectral density estimates were averaged across all artifact-free segments to obtain mean power spectra per channel.

### Procedure

First, written informed consent was obtained from all participants, and a screening questionnaire was completed to identify any contraindications for the stimulation and ensure that all the inclusion criteria were met. Second, participants completed the counting span task to assess their working memory capacity and filled out questionnaires regarding their current state, habits, and demographic information while the electrode cap was equipped. Next, the learning phase of the PALT task followed. Subsequently, the eye-tracking system was calibrated. Then, resting-state EEG was recorded for 5 minutes (the first 2.5 minutes in an eyes-open condition and 2.5 minutes with eyes closed). Following the recording, participants started five practice blocks of the ASRT task, during which stimuli appeared in a completely random order. No statistical learning is expected during these blocks due to the random nature of stimulus presentation. During the random blocks, participants did not receive tACS stimulation. tACS stimulation began when participants started the ASRT task (***tACS session***), which overall consisted of twenty blocks, corresponding to four 5-block epochs (epochs 1-4, Figure 1, Panel D). Participants first completed epochs 1-2 of the ASRT task, and then completed a retrieval phase for the PALT (***tACS retrieval***). The experiment then continued with the remaining ten blocks of the ASRT task (epochs 3-4). Then, another 5-minute-long resting-state EEG recording was conducted post-stimulation. Following the recording, there was a 15-minute break allowing participants to rest. After the break, the eye-tracking device was calibrated again, and participants completed another round of fifteen blocks of ASRT task (***post-tACS session***) with the following structure: first five blocks with the original sequence (epoch 5), followed by five blocks with a previously not practiced, partially overlapping sequence that included both new triplets and triplets according to the original sequence (i.e., two of the four pattern elements remained the same, while the other two were switched; epoch 6), ending with the final five blocks containing solely the original sequence again (epoch 7). Participants were unaware of changes between the sequences. Finally, participants repeated a retrieval phase for the PALT (***post-tACS retrieval***). An experimental session lasted for approximately 3.5 hours (see Figure 1, Panel D).

### Data preparation and statistical analysis

Prior to analysis, trials were categorized based on whether they served as the last element of a high-probability or a low-probability triplet in a sliding window manner. It has been shown that preexisting response tendencies exist for specific elements, which may introduce a bias to the RT assessment; thus, trills (e.g., 1-2-1) were excluded (Howard et al. 2004). Because the subsequent presentation of the target stimulus at the same location would result in 0-ms-long RTs that would introduce a bias in the estimates, we also excluded repetitions (e.g., 2-2). Additionally, to control for extreme outliers, trials with an RT outside three standard deviations from the individual mean RT were excluded. Overall, the complete data preparation resulted in the removal of 28.75% of all trials. During data analysis, each five blocks was further nested into epochs.

Statistical analyses and data visualization were performed using custom Python scripts (version 3.13.3) and RStudio (version 2024.12.1.563; Posit team 2025). Linear Mixed Models (LMMs) were employed using the *lme4* package (version 1.1.36) for reaction time analysis. A generalized linear mixed-effects model (GLMM) was conducted in the presence of various measures of pre-stimulus gaze direction changes. Estimated marginal means were computed with the *emmeans* R package. Post hoc comparisons were corrected for multiple comparisons using the Holm–Bonferroni method. An alpha level of 0.05 was applied to all analyses. Figures were created with the *ggplot2* package. We carried out an iterative random effects structure simplification, starting with the maximal random-effects structure, which included random effects for all variables and allowed correlations between them. We then reduced this structure to achieve convergence by removing the correlations between random slopes, or the random slopes themselves, if necessary. The numerical fixed factors were mean-centered. For each model, we report the full model structure in the Results section. Importantly, differences in statistical learning are indicated by the Triplet Type main effect and interactions involving this factor. Main effects and interactions without the Triplet Type can be interpreted as differences in non-specific changes of visuomotor performance.

To analyze PALT scores, we conducted mixed-design ANOVAs using JASP (version 0.19.3; JASP Team 2024), with the given score (associate learning, item, or recollection) as the dependent variable, while time (during stimulation, post-stimulation) was a within-subject factor, and group (sham, synchronization) was a between-subject factor.

We also performed mixed-design ANOVAs to analyze the brain oscillatory changes at 6 Hz over the region of interest that included the following electrode sites: Fp1, Fp2, F3, F4, and Fz. In this case, the power of 6 Hz activity in dB over the region of interest was entered as the dependent variable, time (pre-stimulation, post-stimulation) was a within-subject factor, and group (sham, synchronization) was entered as a between-subject factor.

## Results

The groups were comparable regarding age (*M*_sham_ = 21.31, *SD*_sham_ = 0.47; *M*_synchronization_ = 21.57, *SD*_synchronization_ = 0.55, *p* = 0.727), sex (*n*_sham_ = 6 male, *n*_synchronization_ = 9 male, *p* = 0.069), handedness (*n*_sham_ = 1 left-handed, 1 ambidextrous; *n*_synchronization_ = 1 left-handed, *p* = 0.773), and education in years (*M*_sham_ = 14.50, *SD*_sham_ = 2.06; *M*_synchronization_ = 14.97, *SD*_synchronization_ = 2.06, *p* = 0.507).

Statistical learning during the tACS session was evident in both groups in terms of reaction time (see Figure 2) as well as learning-dependent anticipations; however, no difference was found between the two experimental groups (see Supplementary Results for statistical analysis). None of the participants reported explicit knowledge of the sequential structure of the ASRT task.

**Figure 2.**
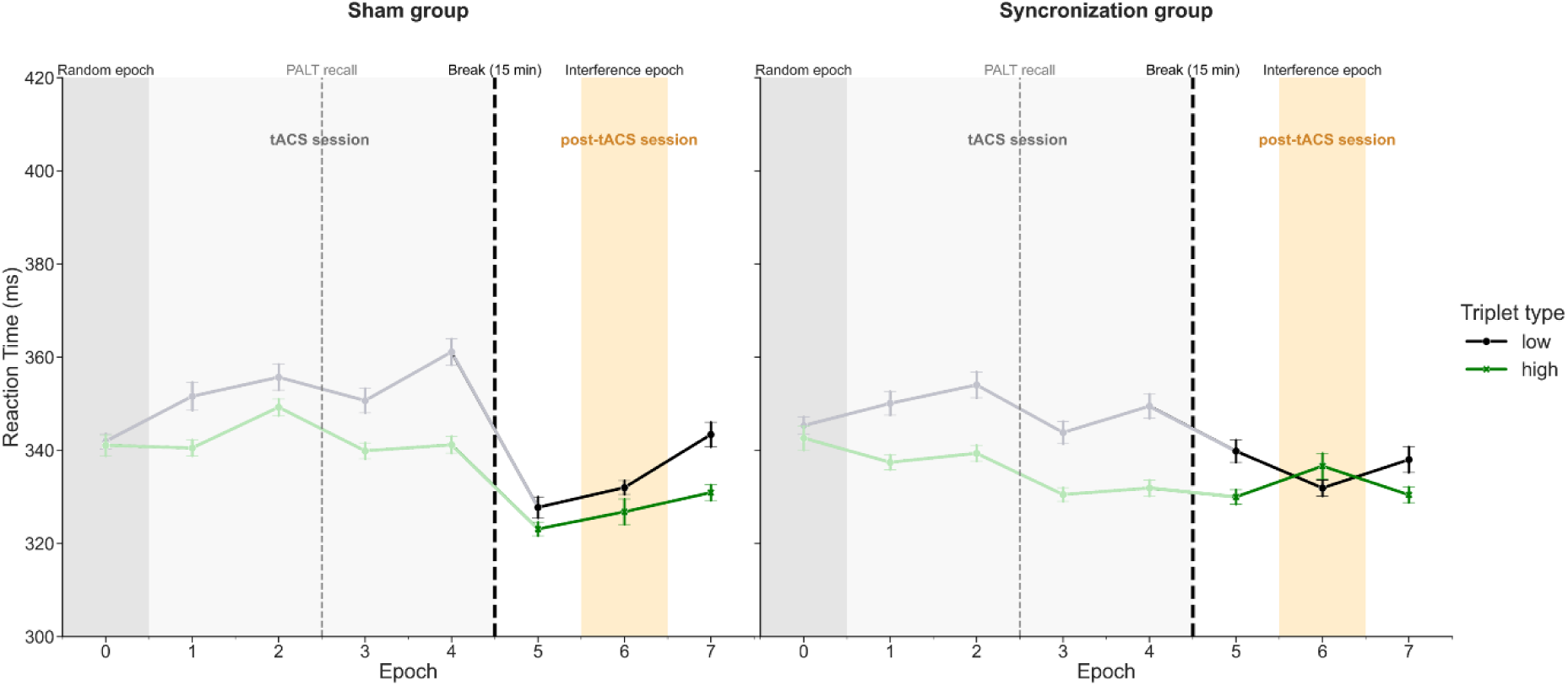
The trajectory of reaction times as a function of high-probability (green line) and low-probability (black line) triplets throughout the experiment. Note that stimuli were presented randomly in Epoch 0 (marked as dark grey), and participants were presented with a new sequence at Epoch 6 (marked by the peach color). The difference between high- and low-probability triplets (coded according to the original sequence) represents statistical learning. In the tACS session, statistical learning was statistically significant by Epoch 4 in both groups (see Supplementary Results). Error bars show the standard error of the mean.

### Are the prolonged effects of theta synchronization evident in statistical learning (reaction time)?

To specifically assess consolidation and if tACS had any prolonged effects, an LMM regressing reaction time (outcome variable) on Triplet Type (high-probability vs low-probability, according to the original sequence, entered as a factor), Epoch (5-7, entered as a factor), and Group (Sham vs Synchronization, entered as a factor) was employed. The random effect structure included correlated random intercepts and random slopes for Triplet Type per subject number. The main effect of Triplet Type was once again significant, suggesting statistical learning with faster RTs for high-probability trials compared to low-probability trials (*F*_(1, 56.77)_ = 7.54, *p* = 0.008). The Epoch main effect showed a change in visuomotor performance in the interference epoch (i.e., epoch 6), indicated by slower RTs (*F*_(2, 40833.77)_ = 7.20, *p* < 0.001). There was a performance change in statistical learning according to the original sequence, as well as indicated by the Triplet Type × Epoch interaction (*F*_(2, 40822.17)_ = 7.17, *p* < 0.001). In the interference epoch, the difference between high-probability and low-probability trials according to the original sequence was not significant, as expected (*b*_epoch 5_ = -7.56, *SE*_epoch 5_ = 2.77, *p*_epoch 5_ = 0.006; *b*_epoch 6_ = 0.31, *SE*_epoch 6_ = 2.78, *p*_epoch 6_ = 0.912; *b*_epoch 7_ = -10.69, *SE*_epoch 7_ = 2.77, *p*_epoch 7_ < 0.001). Moreover, a statistically significant difference was found in the Group × Epoch interaction, indicating that the progress of visuomotor learning differed between the two groups (*F*_(2, 40833.77)_ = 8.17, *p* < 0.001). However, after correcting for multiple comparisons, there was no difference in RTs across the groups at any epoch (*b*_epoch 5_ = -9.03, *SE*_epoch 5_ = 8.84, *p*_epoch 5_ = 0.307; *b*_epoch 6_ = -4.76, *SE*_epoch 6_ = 8.84, *p*_epoch 6_ = 0.591; *b*_epoch 7_ = 2.86, *SE*_epoch 7_ = 8.84, *p*_epoch 7_ = 0.747). We did not find further significant higher-order interactions. Full results are reported in Supplementary Table S2.

### Are the prolonged effects of theta synchronization evident in pre-stimulus gaze directions?

#### 1. The amount of pre-stimulus gaze direction changes from the quadrant of the previous stimulus

We fitted a GLMM with a binomial error distribution to examine pre-stimulus gaze direction changes (dichotomous outcome measure). The model included Epoch (5–7, entered as a factor) and Group (Sham vs. Synchronization, entered as a factor) as fixed effects. Random intercepts and correlated random slopes for Epoch were included for each subject.

The model showed a significant effect of Epoch for Epoch 7 (*z* = 3.344, *p* = 0.001), but not for Epoch 6 (*z* = 1.331, *p* = 0.183) compared to Epoch 5, indicating that participants regardless of group assignment were more likely to move their gaze pre-stimulus from the quadrant where the previous stimulus was presented in the last epoch (*b*_epoch5-6_ = -0.344, *SE*_epoch5-6_ = 0.084, *p*_epoch5-6_ = 0.001; *b* _epoch5-7_ = -0.570, *SE* _epoch5-7_ = 0.100, *p*_epoch5-7_ < 0.001). There was a tendency in the sham group compared to the synchronization group to change their gaze direction pre-stimulus (*z* = -1.948, *p* = 0.051). Moreover, as indicated by the Epoch × Group interaction, there was a difference between groups in the amount of pre-stimulus gaze direction changes at Epoch 6 (*z* = 2.574, *p* = 0.010) and Epoch 7 (*z* = 1.682, *p* = 0.093). Post hoc test revealed that the sham group had more pre-stimulus gaze direction changes compared to the synchronization group, but the difference was not statistically significant at any epoch after correcting for multiple comparisons (*b*_epoch 5_ = 0.813, *SE*_epoch 5_ = 0.417, *p*_epoch 5_ = 0.051; *b*_epoch 6_ = 0.394, *SE*_epoch 6_ = 0.367, *p*_epoch 6_ = 0.283; *b*_epoch 7_ = 0.483, *SE*_epoch 7_ = 0.326, *p*_epoch 7_ = 0.138). Higher-order interactions were not statistically significant. Full results are reported in Supplementary Table S3.

#### 2. Learning-dependent anticipation ratio

An LMM regressing reaction time (outcome variable) on Triplet Type (high-probability vs low-probability, according to the original sequence, entered as a factor), Epoch (5-7, entered as a factor), and Group (Sham vs Synchronization, entered as a factor) was conducted to assess the online effects of tACS on statistical learning. The random effect structure included random intercepts per subject number. The main effect of Epoch (*F*_(2, 349.33)_ = 6.52, *p* =0.002) revealed that the amount of learning-dependent anticipations dropped during the interference epoch, at Epoch 6 compared to Epoch 5 (b_epoch 5-6_ = 9.50, SE_epoch 5-6_ = 2.82, *p*_epoch 5-6_ = 0.002), and increased again in Epoch 7 (*b*_epoch 5-7_ = 2.97, *SE*_epoch 5-7_ = 3.21, *p*_epoch 5-7_ = 0.626). The Epoch × Group interaction (*F*_(2, 349.33)_ = 0.40, *p* =0.668) was not significant; full results are reported in Supplementary Table S4.

#### 3. Learning-dependent anticipations

A GLMM with a binomial error distribution was fitted, where trial type (trials with learning-dependent anticipation, other trials) was entered as the outcome measure. The model included Epoch (5–7, entered as a factor) and Group (Sham vs. Synchronization, entered as a factor) as fixed effects. Random intercepts and correlated random slopes for Epoch were included for each subject. The main effect of Epoch revealed that the amount of learning-dependent anticipations decreased during the interference epoch at Epoch 6 (*z* = -2.58, *p* = 0.040) and increased by Epoch 7 (*z* = 4.044, *p* < 0.001) compared to Epoch 5. The main effect of Group (*z* = 3.60, *p* = 0.100), and the higher-order interactions were non-significant. Full results are reported in Supplementary Table S5.

### Do prolonged effects of theta synchronization affect old and new knowledge during the interference epoch?

As described previously, to examine how participants reacted to knowledge acquired during the stimulation and new knowledge learnt following the stimulation, we categorized stimuli into four ***Information Types*** (HH, HL, LH, and LL) in the interference epoch (i.e., Epoch 6). In this analysis, LL is considered a reference level, as it shows performance to low-probability triples according to both the old and new sequences.

#### 1. Reaction time

An LMM regressing reaction time (outcome variable) on Information Type (HH, HL, LH, and LL, entered as a factor), and Group (Sham vs Synchronization, entered as a factor) was employed. The random effect structure included random intercepts per subject number. There was a significant interaction between Information Type and Group (*F*_(3, 20916.14)_ = 2.69, *p* = 0.045). Post hoc tests, however, revealed no difference between the RT for any information type compared to the reference level of LL (*b*_LH_ = 3.66, *SE*_LH_ = 8.90, *p*_LH_ = 1.000; *b*_HL_ = -1.87, *SE*_HL_ = 5.62, *p*_HL_ = 1.000; *b*_HH_ = 17.48, *SE*_HH_ = 7.51, *p*_HH_ = 0.100). Full results are reported in Supplementary Table S6.

#### 2. Learning-dependent anticipations

We regressed the amount of learning-dependent anticipations in all trials in Epoch 6 (interference epoch) using a GLMM with a binomial error distribution on Information Type (HH, HL, LH, and LL, entered as a factor) and Group (Sham vs Synchronization, entered as a factor). The random effect structure included random intercepts per subject number. The percentage of learning-dependent anticipations according to Information Types tendentially differed from the reference level of LL for both LH (*z* = -1.896, *p* = 0.058), HL (*z* = -1.854, *p* = 0.064), and significantly differed in the case of HH (*z* = 5.068, *p* < 0.001). The main effect of Group was not significant (*z* = -0.493, *p* = 0.622). Importantly, we found that learning-dependent anticipations differed across Groups and Information Types. As indicated by the Information Type × Epoch interaction, compared to the reference level of LL in the Sham group, the learning-dependent anticipations significantly differed in the Synchronization group LH (*z* = 2.753, *p* = 0.006), HH (*z* = -3.650, *p* < 0.001). At the same time, we did not find a difference in HL (*z* = 1.880, *p* = 0.060). Post hoc analysis using LL as a reference level revealed that participants had comparable amounts of learning-dependent anticipations in the HL condition (*b*_HL_ = 0.382, *SE*_HL_ = 0.203, *p*_HL_ = 0.120). This suggests that participants could disengage equally well from the previously high-probability triplets once they became low-probability. Importantly, the amount of learning-dependent anticipations differed across the two groups regarding the LL-LH (*b*_LH_ = 0.346, *SE*_LH_ = 0.126, *p*_LH_ = 0.017). Participants in the Sham group, compared to those in the Synchronization group, exhibited a lower propensity for learning-dependent anticipations in the case of triplets that newly became high-probability, suggesting more efficient predictions to newly learned high-probability triplets for the Synchronization group. Additionally, the LL-HH difference also differed across the two groups (*b*_HH_ = -0.603, *SE*_HH_ = 0.165, *p*_HH_ = 0.001). Overall, participants in the Sham group had more learning-dependent anticipations than participants in the Synchronization group for HH stimuli, indicating more stable performance for already acquired knowledge in the Sham group (Figure 3). Moreover, participants in the Synchronization group seemed to disengage from the previously learned sequence easily. They showed more learning-dependent anticipations for LH stimuli, while the Sham group showed elevated learning-dependent anticipation for HH stimuli. Full results are reported in Supplementary Table S7.

**Figure 3.**
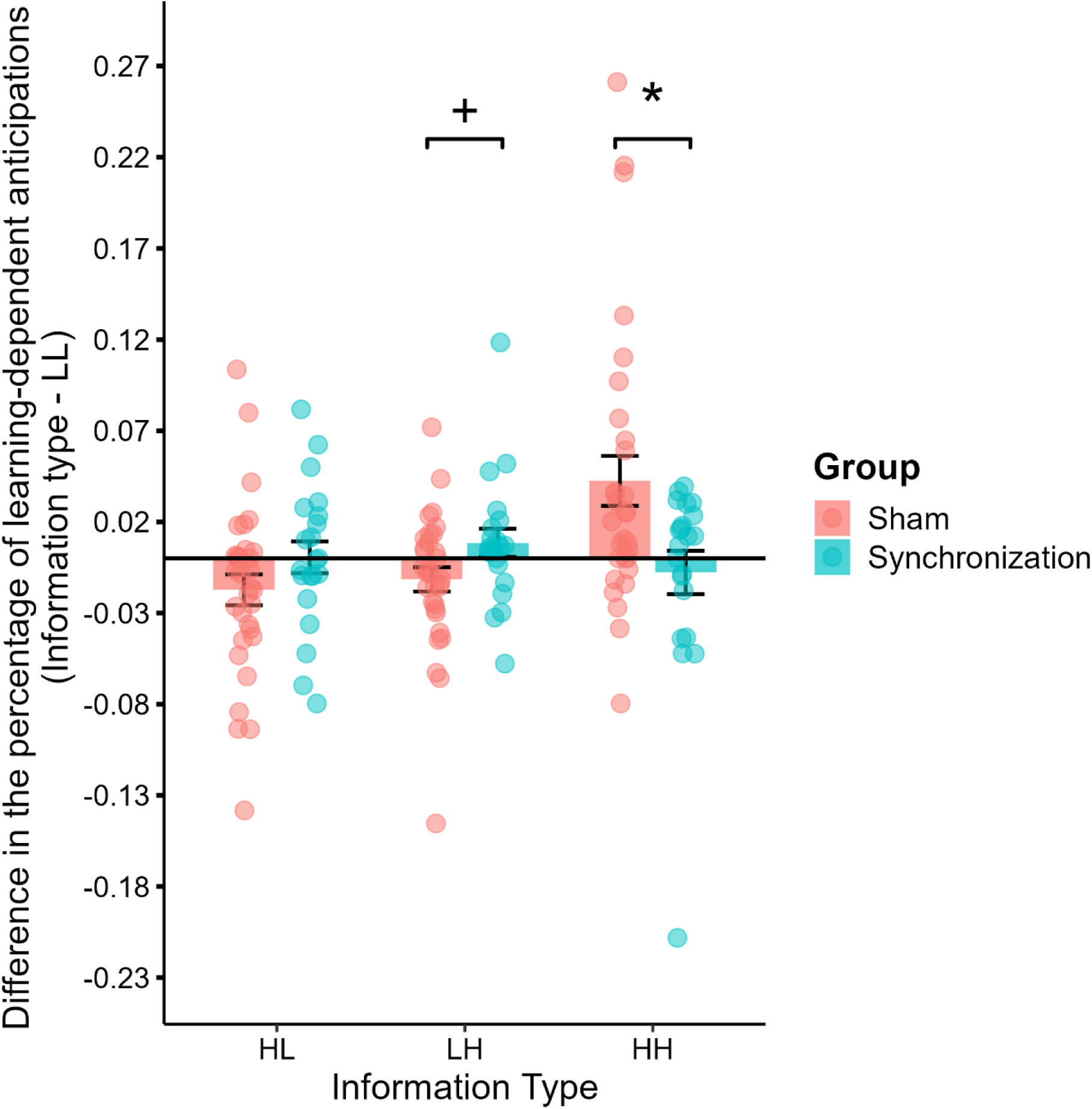
The percentage of learning-dependent anticipations controlled for the reference level of low-low (LL) as a function of group and information type in Epoch 6. The Sham group has significantly more learning-dependent anticipations for HH than the Synchronization group. The Synchronization group had more learning-dependent anticipations for LH than the Sham group. *Note. HL: high-low, LH: low-high, HH: high-high*

### Are the prolonged effects of theta synchronization evident in the likelihood of pre-stimulus gaze direction types?

Finally, we employed a more nuanced analysis of pre-stimulus gaze directions that categorized all trials into four categories. In this analysis, we integrated both the probability structure of the task, by weighing each type of anticipatory eye movement according to its chance-level probability, and the actual accuracy of the anticipations. An LMM was employed to regress reaction time (outcome variable) on Pre-stimulus Gaze Direction Type (LDC, LDE, NLDC, and NLDE, entered as a factor), Epoch (5-7, entered as a factor), and Group (Sham vs Synchronization, entered as a factor). We found a significant Pre-stimulus Gaze Direction Type × Group interaction (*F*_(3, 561)_ = 5.94, *p* < 0.001). The Sham group showed significantly more LDEs than the Synchronization group (*b* = 0.096, *SE* = 0.023, *p* < 0.001), indicating that the Synchronization group made fewer errors by anticipating high-probability triplets in the post-tACS session (Figure 4). Combined with the fewer anticipations for HH trials, this result suggests a more flexible processing in the Synchronization group, which is marked by faster adaptation to the new sequence (LH gain) and fewer anticipations for the previously acquired sequence once the probabilities have changed with fewer learning-dependent errors. Full results are reported in Supplementary Table S8.

**Figure 4.**
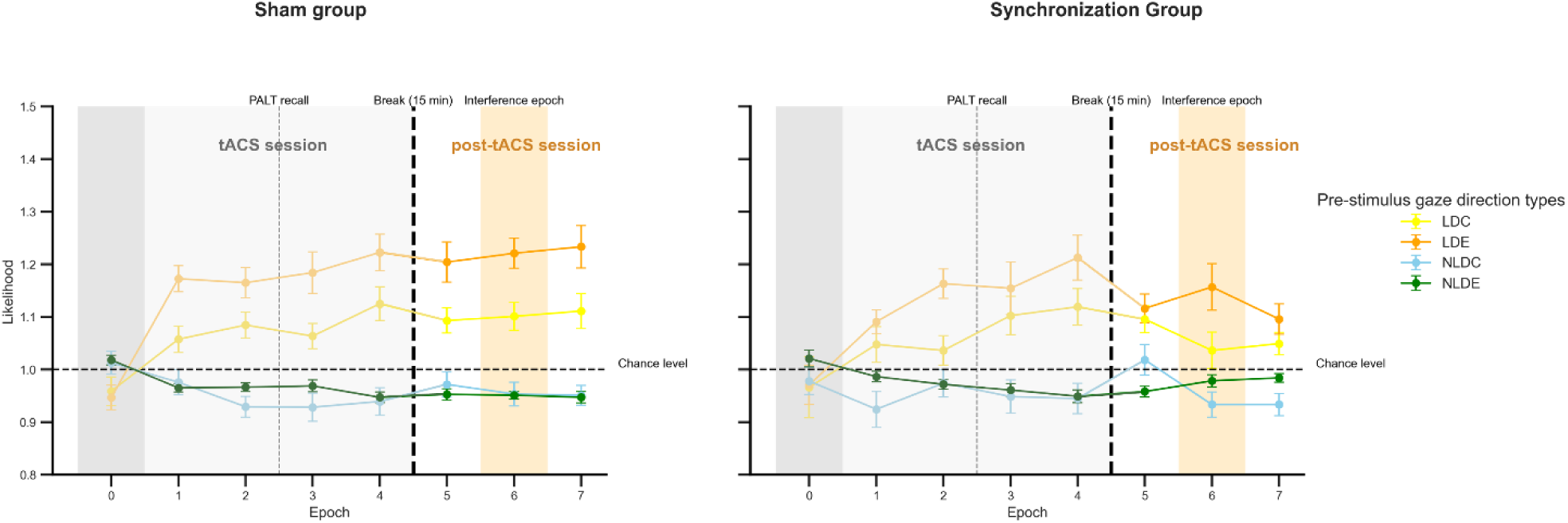
The likelihood of pre-stimulus gaze direction types per epoch per group. There was a significant difference in the likelihood of LDEs (orange line) between the two groups, while LDC (blue line), NLDC (green line), and NLDE (red line) did not differ between groups. The chance level is marked with a dashed line. Error bars show the standard error of the mean. *Note: LDC: Learning-dependent correct, LDE: Learning-dependent error, NLDC: Not-learning-dependent correct, NLDE: Not-learning-dependent error*

### Are the prolonged effects of theta synchronization evident in 6 Hz power on the resting-state EEG?

The ANOVA revealed a non-significant main effect of Time (*F*(1, 58) = 2.351, *p* = 0.131, η_p_² = 0.039), Group (*F*(1, 58) = 0.014, *p* = 0. 906, η_p_² < 0.001), and a non-significant Time × Group interaction (*F*(1, 58) = 0.057, *p* = 0.813, η_p_² < 0.001) for the eyes-open resting-state EEG. Similarly, a non-significant main effect of Time (*F*(1, 58) = 0.398, *p* = 0.725, η_p_² = 0.012), Group (*F*(1, 58) = 0.704, *p* = 0.405, η_p_² = 0.012), and a non-significant Time × Group interaction (*F*(1, 58) = 0.521, *p* = 0.473, η_p_² = 0.009) was found for the eyes-closed resting-state EEG.

### Are the prolonged effects of theta synchronization evident in a control declarative memory task?

The ANOVA for the associate learning score revealed a significant main effect of Time (*F*(1, 58) = 5.012, *p* = 0.029, η_p_² = 0.080). Participants, regardless of group assignment, had a higher associate learning score in the post-tACS session compared to the tACS session (*M*_session 1 - session 2_ = -0.133, *SE*_session 1 - session 2_ = 0.059, *p* = 0.029). The Group main effect (*F*_(1, 58)_ = 0.290, *p* = 0.592, η_p_^2^ = 0.005), as well as the Time × Group interaction (*F*_(1, 58)_ = 1.767, *p* = 0.189, η_p_^2^ = 0.030) were not significant. Similarly, no change was revealed using the item and recollection scores. For detailed results, see Supplementary Results.

## Discussion

In the present study, we aimed to go beyond assessing the online effects of frontal midline theta synchronization and measured the delayed aftereffects of the stimulation on both the retention of previously acquired probabilistic sequences and the learning of a new, partially overlapping sequence structure. Utilizing a sensitive eye-tracking paradigm allowed us to quantify predictive processing through pre-stimulus gaze directions. Critically, broad performance metrics, such as reaction times and overall anticipatory behaviors, did not differ significantly between the synchronization and control groups. However, our nuanced analyses, including the use of pre-stimulus gaze direction types as well as the detailed investigation of how old and new knowledge were affected by tACS in the interference epoch, revealed a subtle yet significant enhancement in cognitive flexibility among participants who received active stimulation, all without mitigating the previously acquired statistical regularities. Our results suggest that frontal midline theta synchronization enhances the adaptation of predictive processes, specifically, highlighting its potential to facilitate neural plasticity in complex learning contexts.

The lengthy updating of prior mental representations in the sham group is consistent with previous findings. Kóbor et al. (2020) have shown that the knowledge of an already learned or over-practiced probabilistic sequence may persist, even after changing the transitional probabilities by providing completely unstructured (random) blocks (Kóbor et al. 2020). A similar tendency to adhere to prior knowledge has also been described in an eye-tracking study using the same task, both for reaction time and learning-dependent anticipations when a novel, interfering structure was introduced (Zolnai et al. 2022). Although distinct representations for old and new sequences can coexist without mutual inhibition, the stabilization of a novel sequence structure typically requires extended practice over multiple epochs (Horváth et al. 2022; Takács et al. 2025). In contrast, the present study demonstrates that frontal synchronization in theta frequency expedited this adaptive process, accelerating the updating of internal probabilistic models in the synchronization group.

Enhanced cognitive flexibility, i.e., more effective updating of mental representations, was demonstrated in the synchronization group by more learning-dependent anticipations corresponding to the new regularity during the interference epoch compared to the sham group. Additionally, the synchronization group had significantly fewer learning-dependent anticipations that corresponded to the old regularity, as well as fewer learning-dependent errors in reference to the sham group. This is in line with research showing that boosting frontal midline theta activity—using either non-invasive brain stimulation (Lehr et al. 2019; Lu et al. 2024) or neurofeedback (Enriquez-Geppert et al. 2014; Smit et al. 2023)—can increase cognitive flexibility and adaptive behavior. Together, these findings indicate that synchronization of frontal midline theta enabled more flexible switching from the previously acquired statistical probabilities to partially new probabilistic sequences.

Several underlying cognitive processes can explain the observed differences between the sham and synchronization groups. One possibility is that boosting the frontal midline theta shifted participants in the synchronization group toward a prefrontal cortex-driven, model-based processing strategy, as opposed to the more model-free and implicit processing seen in the sham group. While none of our participants reported explicit knowledge of the sequence, this shift in cognitive strategy may have facilitated the executive functions, resulting in the more flexible updating of predictive processes and better inhibition of knowledge of the prior sequence. In a similar fashion, explicit cues about new regularities were found to promote the rewiring of representations compared to when the learning of the novel sequence happened implicitly (Szegedi-Hallgató et al. 2017). A model-based strategy does not necessarily equate to full explicit awareness; however, it can be speculated that explicit cues in the task may have facilitated model-based processing (Castro-Rodrigues et al. 2022). Nevertheless, this suggests that the acquisition and the rewiring of statistical regularities rely on distinct cognitive mechanisms, the latter supported by prefrontal cortex-related activity.

Another speculation is that, instead of learning simply to distinguish high- and low-probability triplets, theta synchronization may allow participants to acquire a more abstract representation of the task structure that was valid for both the old and new sequence. Piling evidence suggests that statistical learning relies on concomitant coding on several levels, including a stimulus (S-cluster), a response (R-cluster), and a more abstract stimulus-response or cognitive level (C-cluster) (Takács et al. 2021; Vékony et al. 2023). It is possible that our intervention boosted the development of the latter, i.e., high-level structure representations that incorporate context probability in addition to general sensorimotor contingencies (Takács et al. 2025). If this is the case, participants in the synchronization group may have developed a more general model of the task, recognizing the alternating structure, allowing them to surpass the need to inhibit or update the previous sequence. This would mean that frontal midline theta synchronization allowed participants to flexibly change how their existing memories are expressed by quickly adapting to a novel context without having to inhibit the previously acquired knowledge.

Our findings may also be interpreted in terms of frontal midline theta synchronization– induced modulation of error monitoring. Errors typically elicit adaptive adjustments such as increased caution and enhanced attentional engagement with task stimuli, processes consistently linked to frontal midline theta oscillations (Cavanagh and Shackman 2015; Valadez and Simons 2018; Beldzik et al. 2022). Indeed, error-related frontal theta has been suggested to reflect the dynamic adjustment of cognitive control during performance monitoring (Cavanagh and Shackman 2015). It is therefore plausible that stimulation in the synchronization group facilitated a shift toward a more cautious strategy. This is also in line with the fewer learning-dependent errors in the synchronization group, which may indicate an elevated sensitivity to errors. However, complementary evidence suggests that automatic error detection and conscious error evaluation are independent of sequence learning and retrieval (Horváth et al. 2021).Thus, while it is possible that error processing played a part in our findings, we expect other mechanisms to also contribute to the effects we found.

One may argue that our results were rather methodological byproducts for two reasons. First, the question could be raised if the synchronization group could develop stable statistical knowledge or could not access the previously acquired structural information in the post-tACS session; however, our results suggested stable acquisition of the original sequence and no difference between the two groups at the beginning or end of the post-tACS session, where the original sequence was re-presented. This latter suggests that theta synchronization did not come at the cost of the knowledge of the original sequence; thus, it can be considered adaptive. Likewise, the question may arise whether the stimulation delivered using the given montage could reach the frontal eye field and alter the quantity or quality of the eye movements regardless of the task. This could be the case as the frontal eye field shows a correlation with microsaccade responses (Hsu et al. 2021) and exploratory eye movements (Becker et al. 2015); however, no significant difference was found between the two groups either for the percentage of pre-stimulus gaze direction changes. Thus, changes in the learning-specific metrics we described are not explained by such a difference.

Our finding that frontal midline theta synchronization did not alter statistical learning during the online stimulation phase aligns with the null results of previous reports (Zavecz et al. 2020; Diedrich et al. 2024). This lack of an immediate effect is not unexpected. A growing body of evidence suggests that the neuroplastic consequences of tACS and other non-invasive brain stimulations often manifest as delayed, offline aftereffects rather than as performance changes during the stimulation itself (Ambrus et al. 2020; Grover et al. 2023). However, the question of whether theta frontal midline theta synchronization could similarly enhance the acquisition of a novel sequence if that is presented during the stimulation remains open.

Null results on the episodic memory task, on the other hand, may seem surprising, considering the involvement of frontal midline theta in episodic memory processes (Eschmann et al. 2020; Yeh et al. 2022; Shtoots et al. 2024). There are multiple explanations for this null effect. First, our task was an incidental learning task where participants did not know that two retrieval phases would occur. During retrieval, prefrontal cortex activity was linked to inhibitory and monitoring control functions (Kim 2010; Manenti et al. 2010; Becker et al. 2015). However, these control processes—together with frontal theta activity—may not be as crucial when participants rely on familiarity-based or weaker memory traces (Guderian and Düzel 2005). Second, it is possible that the task we used was not challenging enough to raise the demand for these top-down processes. Importantly, however, this null effect further demonstrates the specificity of the effect of tACS on the updating of statistical regularities.

The fact that resting-state theta power at 6 Hz did not significantly increase over the frontal electrodes after the stimulation compared to the baseline may also seem unexpected; however, it must be highlighted that this result is not unique in the literature. Despite some promising evidence (Debnath et al. 2025), several articles have failed to identify a theta power increase following theta tACS (Wischnewski et al. 2016; Wischnewski and Schutter 2017). Common reasoning for the lack of tACS-induced resting EEG changes includes methodological choices like the electrode montage. It is possible that larger-scale network-level stimulation is required to increase resting-state theta power (Wischnewski et al. 2016; Pahor and Jaušovec 2018). Another explanation is that transcranial electric brain stimulation tends to work in a winner-takes-all manner, strengthening the already ongoing networks (Fertonani and Miniussi 2016). Considering that we provided stimulation during the task only, it is possible that we could have observed an increased frontal theta power in the task-related networks, but not in networks related to resting state, like the default mode network. This also means that our non-significant results with respect to resting-state EEG does not necessarily mean no modulation of neuronal functioning. Additionally, taking into account individual differences in baseline theta power could have also improved the results. Individualized theta tACS has been shown to result in more pronounced effects on resting-state theta power compared to sham stimulation (Aktürk et al. 2022; Zhang et al. 2022). However, there are methodological challenges to consider in deciding how to calculate individual theta (Wischnewski et al. 2016).

To further elucidate the precise mechanisms by which frontal midline theta synchronization impacts statistical learning, future research could incorporate several methodological advancements. For instance, recording EEG during the task or between learning blocks would enable the direct measurement of on-task theta changes, offering deeper insight into the stimulation’s neurophysiological effects. To further investigate the cognitive background of our results and decide if theta tACS facilitates cognitive flexibility or the development of a more abstract knowledge of the probabilistic, it would be interesting to add either more sequences or repeatedly change the sequence for several epochs. Additionally, future studies should test alternative electrode montages. Targeting distinct brain regions or networks, such as the frontoparietal network with high-density tACS, could reveal how the interplay of distinct areas impacts cognitive flexibility, moving beyond the bilateral prefrontal approach used here (Zavecz et al. 2020).

The present study has two primary limitations. First, the final group sizes were smaller than initially planned, which may affect statistical power. Second, a different electrode montage was used for the sham group compared to the active synchronization group. This decision was a deliberate attempt to address the significant challenge of effective participant blinding in tACS research. By using a denser montage, we aimed to elicit more intense scalp sensations in the sham condition, thereby reducing the potential influence of participants’ expectations on the results (Monaghan et al. 2021). While this approach enhances blinding, we acknowledge it is not an ideal control. To mitigate this in the future, studies should consider alternating the sham montage to match the electrode locations of each active stimulation group. Importantly, the sham stimulation itself was delivered for a very brief period, a duration widely believed to be insufficient to induce significant changes in cortical excitability (Dissanayaka et al. 2017) or EEG power (Pahor and Jaušovec 2018; Hosseinian et al. 2021).

In conclusion, our findings provide evidence for the role of frontal theta oscillations in cognitive flexibility and demonstrate that this enhancement also alters the process of implicit statistical learning. Our work advances theoretical understanding by showing that synchronizing frontal midline theta activity enhances the ability to update existing mental models, positioning it as a key signature for the adaptive reorganization of neural representations. This insight was achieved through a novel methodological approach that combined tACS with a sensitive, eye-tracking-based interference paradigm, a framework that provides a window into the dynamics of predictive processing through the analysis of anticipatory eye movements. Importantly, these measures were sensitive enough to reveal subtle effects that may remain undetected when relying solely on reaction times or pre-stimulus gaze shifts.

## Acknowledgements

This work was supported by the French National Grant Agency (ANR-22-CPJ1-0042-01 and ANR-24-CE37-5807) (to D.N.); the National Brain Research Program project NAP2022-I-2/2022 (to D.N.). The authors are grateful to Zsolt Turi for his help and advice during the project designing, and Botond Becser, Dominika Réka Dávid, Márton Németh, and Dóra Osztényi for their work in the data collection.

## Data Availability Statement

All data presented in this article are available at: https://osf.io/pvz2a/?view_only=ee438b042dcf4428b424c8d4ca3bcece.

## Author contributions

AH: Formal analysis, Visualization, Writing – original draft, Writing—review & editing; OP: Conceptualization, Methodology, Data curation, Software, Formal analysis, Investigation, Project administration, Writing – original draft, Writing—review & editing; TV: Conceptualization, Supervision, Writing—review & editing; GC: Formal analysis, Validation, Supervision, Writing—review & editing; DN: Conceptualization, Funding acquisition, Methodology, Supervision, Writing – review & editing

