## Supplementary Material for "Frontal theta synchronization facilitates the updating of statistical regularities, evidenced by predictive eye movements" (from another version of this preprint: version 1, 2025-08-21)

### Supplementary Methods

#### Eye-tracking data quality

Three measures were used to ensure data quality of the eye-tracker ASRT data (Holmqvist et al., 2012). Precision was ensured by calculating the root-mean-square of the sample-to-sample distance (RMS(S2S)), which shows the differences between successive samples within a fixation (i.e., when the participant fixates on the same spot, such as the target stimulus of the ASRT task). The RMS(S2S) was calculated for the last fixation of each epoch. RMS(S2S) scores were identified using the interquartile range (IQR) method, and participants with outlier epochs were excluded from further analysis. Based on the RMS (S2S) score, we identified five outlier epochs of 4 separate participants. Another precision metric, the root-mean-square of the eye-to-eye distance (RMS(E2E)), was also calculated. The RMS(E2E) score indicates the RMS value of the differences in the two eyes' positions for each sample within a fixation window (except for the samples that did not provide valid data for both eyes). Outlier detection was identical to the one used for the RMS(S2S) scores. Based on the RMS (E2E) score, we identified 17 outlier epochs across 10 separate participants. Additionally, we calculated the percentage of data loss per epoch, defined as the number of samples marked as invalid by the eye-tracker for any reason, including blinks, fast head movements, or changes in head position. If the number of outliers exceeded 20% in any epoch, the participant was excluded from the analysis. Based on the data loss rate, 18 outlier epochs of 5 separate participants were found. Notably, there are overlaps within the epoch marked by these data quality measures. Overall, to maintain the reliability and quality of our data, 15 participants were excluded from further analysis (one person from the sham group, nine from the desynchronization group, and five from the synchronization group).

#### Data exclusion

We excluded the ASRT data of four participants due to experimenter error, and two further participants due to corrupted/missing ASRT data from the Sham group. Ten participants' data were excluded from the Desynchronization group due to experimenter error, and one additional participant was excluded due to corrupted/missing ASRT data. Finally, six participants' data were removed from the Synchronization group due to experimenter error, and an additional participant's data were removed due to corrupted/missing ASRT data. The Desynchronization group had more experimenter errors as the montage required the change of electrode locations for the tACS session, which increased the risk of experimenter errors.

#### Supplementary Results

##### Does theta synchronization affect statistical learning in terms of reaction time?

An LMM regressing reaction time (outcome variable) on Triplet Type (high probability vs low probability, entered as a factor), Epoch (1-4, entered as a factor), and Group (Sham vs Synchronization, entered as a factor) was conducted to assess the online effects of tACS on statistical learning. The random effect structure included random intercepts and random slopes for Epoch per subject number. The model showed a main effect of Triplet Type, with faster RTs for high-probability trials than for low-probability trials ($F_{(1, 54316.32)} = 136.74$, $p < 0.001$). Reaction times decreased as the task progressed, indicating visuomotor performance change as evidenced by the Epoch main effect ($F_{(3, 58.56)} = 4.56$, $p = 0.006$). Statistical learning over the progress of the ASRT task was confirmed by a significant Triplet Type × Epoch interaction ($F_{(3, 54308.74)} = 3.17$, $p = 0.023$). As the task progressed, the RT difference between high-probability and low-probability triplets increased ($b_{epoch\ 1} = -11.61$, $SE_{epoch\ 1} = 2.27$, $p_{epoch\ 1} < 0.001$; $b_{epoch\ 2} = -9.88$, $SE_{epoch\ 2} = 2.27$, $p_{epoch\ 2} < 0.001$;$b_{epoch\ 3} = -12.47$, $SE_{epoch\ 3} = 2.26$, $p_{epoch\ 3} < 0.001$; $b_{epoch\ 4} = -19.11$, $SE_{epoch\ 4} = 2.27$, $p_{epoch\ 4} < 0.001$). No further significant higher-level interactions were observed, indicating that RTs and statistical learning were not affected by tACS during the stimulation phase.

| | **RT** | | | | |
|---|---|---|---|---|---|
| *Terms* | *b* | *SE b* | *95% CI* | *t* | *p* |
| (Intercept) | 345.47 | 4.57 | 336.52 – 354.42 | 75.65 | **<0.001** |
| Triplet Type | -6.63 | 0.57 | -7.74 – -5.52 | -11.69 | **<0.001** |
| Epoch 2 | -0.67 | 2.03 | -4.66 – 3.32 | -0.33 | 0.742 |

| | | | | | |
|---|---|---|---|---|---|
| Epoch 3 | 4.43 | 2.02 | 0.48 – 8.38 | 2.20 | **0.028** |
| Epoch 4 | -4.29 | 1.26 | -6.77 – -1.82 | -3.40 | **0.001** |
| Group | 3.20 | 4.57 | -5.75 – 12.15 | 0.70 | 0.484 |
| Triplet Type × Epoch 2 | 0.83 | 0.98 | -1.10 – 2.76 | 0.84 | 0.400 |
| Triplet Type × Epoch 3 | 1.69 | 0.98 | -0.23 – 3.62 | 1.72 | 0.085 |
| Triplet Type × Epoch 4 | 0.40 | 0.98 | -1.52 – 2.32 | 0.41 | 0.685 |
| Triplet Type × Group | 0.77 | 0.57 | -0.34 – 1.88 | 1.35 | 0.176 |
| Epoch 2 × Group | -2.13 | 2.03 | -6.12 – 1.86 | -1.05 | 0.295 |
| Epoch 3 × Group | -0.63 | 2.02 | -4.58 – 3.33 | -0.31 | 0.757 |
| Epoch 4 × Group | 0.66 | 1.26 | -1.81 – 3.13 | 0.53 | 0.599 |
| Triplet Type × Epoch 2 × Group | -0.52 | 0.98 | -2.44 – 1.41 | -0.52 | 0.600 |
| Triplet Type × Epoch 3 × Group | 1.77 | 0.98 | -0.16 – 3.69 | 1.80 | 0.072 |
| Triplet Type × Epoch 4 × Group | 0.05 | 0.98 | -1.87 – 1.97 | 0.05 | 0.960 |
| **Random Effects** | | | | | |
| $\sigma^2$ | 13661.68 | | | | |
| $\tau_{00}$ subject number | 1041.25 | | | | |
| $\tau_{11}$ subject number × Epoch2 | 160.90 | | | | |
| $\tau_{11}$ subject number × Epoch3 | 157.07 | | | | |
| $\tau_{11}$ subject number × Epoch4 | 31.86 | | | | |
| $\rho_{01}$ | -0.13 | | | | |
| | 0.42 | | | | |
| | -0.59 | | | | |
| ICC | 0.08 | | | | |
| N | 53 | | | | |
| Observations | 54482 | | | | |
| Marginal $R^2$ / Conditional $R^2$ | 0.004 / 0.082 | | | | |

#### Does theta synchronization affect pre-stimulus gaze directions?

1. **The amount of pre-stimulus gaze direction changes from the quadrant of the previous stimulus**

We fitted a generalized linear mixed-effects model (GLMM) with a binomial error distribution to examine pre-stimulus gaze direction changes (dichotomous outcome measure). The model included Epoch (1–4, entered as a factor) and Group (Sham vs. Synchronization, entered as a factor) as fixed effects. Random intercepts and random slopes for Epoch were included for each subject.

Participants in the sham group showed a weak but higher overall tendency to move their gaze out of the quadrant where the last stimulus appeared before the next stimulus was presented ($z = -1.825$, $p = 0.068$) during the stimulation phase. No other main effects or interactions were significant.

| *Terms* | *b* | *SE b* | *95% CI* | *z* | *p* |
|---|---|---|---|---|---|
| (Intercept) | -1.2944 | 0.2322 | -1.7495 – -0.8393 | -5.5749 | **<0.001** |
| Epoch 2 | -0.0161 | 0.1081 | -0.2281 – 0.1958 | -0.1491 | 0.881 |
| Epoch 3 | -0.1346 | 0.1342 | -0.3976 – 0.1283 | -1.0035 | 0.316 |
| Epoch 4 | -0.0319 | 0.1342 | -0.2950 – 0.2312 | -0.2374 | 0.812 |
| Group | -0.6690 | 0.3665 | -1.3872 – 0.0493 | -1.8254 | 0.068 |
| Epoch 2 × Group | 0.1899 | 0.1739 | -0.1508 – 0.5307 | 1.0924 | 0.275 |
| Epoch 3 × Group | 0.0159 | 0.2153 | -0.4060 – 0.4378 | 0.0739 | 0.941 |
| Epoch 4 × Group | 0.1718 | 0.2144 | -0.2485 – 0.5920 | 0.8011 | 0.423 |
| **Random Effects** | | | | | |
| $\sigma^2$ | 3.29 | | | | |

| | |
|---|---|
| $\tau_{00}$ subject number | 1.75 |
| $\tau_{11}$ subject number.epoch2 | 0.32 |
| $\tau_{11}$ subject number.epoch3 | 0.51 |
| $\tau_{11}$ subject number.epoch4 | 0.52 |
| $\rho_{01}$ | -0.51 |
| | -0.10 |
| | -0.37 |
| ICC | 0.34 |
| N | 53 |
| Observations | 54482 |
| Marginal $R^2$ / Conditional $R^2$ | 0.017 / 0.349 |

**2. Learning dependent anticipation ratio**

An LMM regressing epoch-wise learning-dependent anticipation ratio (outcome variable) on Triplet Type (high-probability vs low-probability, entered as a factor), Epoch (1-4, entered as a factor), and Group (Sham vs Synchronization, entered as a factor) was conducted. The model, including only the random intercepts per subject number in the random effect structure, was singular. Thus, we ran a likelihood ratio test to assess whether the model outperformed the null model that only included the fixed effects. The full model did not significantly improve fit over the null model ($\chi^2_{(1)} < 0.001$, $p = 0.999$). Additionally, we further ran a GLMM using the trial-wise learning-dependent anticipations (see results below).

**3. Learning-dependent anticipations**

A GLMM with a binomial error distribution was fitted, where learning-dependent anticipations were entered as the outcome measure. The model included Epoch (1–4, entered as a factor) and Group (Sham vs. Synchronization, entered as a factor) as fixed effects. Random intercepts and random slopes for Epoch were included for each subject.

Compared to Epoch 1, participants had significantly more learning-dependent anticipations at Epoch 4 ($z = 3.040$, $p = 0.002$). Similarly to pre-stimulus gaze direction changes, there was a tendency toward more learning-dependent anticipations in the Sham group compared to the synchronization group ($z$ = -1.824, $p$ = 0.068). However, no further significant results were found.

| *Terms* | *b* | *SE b* | *95% CI* | *z* | *p* |
|---|---|---|---|---|---|
| (Intercept) | -2.4957 | 0.1882 | -2.8647 – -2.1268 | -13.2584 | **<0.001** |
| Epoch 2 | -0.0111 | 0.0532 | -0.1153 – 0.0931 | -0.2087 | 0.835 |
| Epoch 3 | -0.0052 | 0.0531 | -0.1093 – 0.0989 | -0.0975 | 0.922 |
| Epoch 4 | 0.1574 | 0.0518 | 0.0559 – 0.2589 | 3.0403 | **0.002** |
| Group | -0.5507 | 0.3019 | -1.1424 – 0.0409 | -1.8244 | 0.068 |
| Epoch 2 × Group | 0.1005 | 0.0920 | -0.0798 – 0.2808 | 1.0921 | 0.275 |
| Epoch 3 × Group | 0.0116 | 0.0928 | -0.1703 – 0.1935 | 0.1253 | 0.900 |
| Epoch 4 × Group | 0.0096 | 0.0904 | -0.1676 – 0.1867 | 0.1057 | 0.916 |
| **Random Effects** | | | | | |
| $\sigma^2$ | 3.29 | | | | |
| $\tau_{00}$ subject number | 1.10 | | | | |
| ICC | 0.25 | | | | |
| N | 53 | | | | |
| Observations | 54482 | | | | |
| Marginal $R^2$ / Conditional $R^2$ | 0.016 / 0.261 | | | | |

**Does theta synchronization affect pre-stimulus gaze direction types?**

An LMM was employed to regress reaction time (outcome variable) on pre-stimulus gaze direction types (LDC, LDE, NLDC, and NLDE, entered as a factor), Epoch (1-4, entered as a factor), and Group (Sham vs Synchronization, entered as a factor). The random effect structure included random intercepts per subject number. There were no differences between the two Groups.

| *Terms* | *b* | *SE b* | *95% CI* | *t* | *p* |
|---|---|---|---|---|---|
| (Intercept) | 1.04 | 0.01 | 1.03 – 1.05 | 179.75 | **<0.001** |
| Pre-stimulus gaze direction type (LDE) | 0.04 | 0.01 | 0.02 – 0.06 | 4.75 | **<0.001** |
| Pre-stimulus gaze direction type (NLDC) | 0.13 | 0.01 | 0.11 – 0.15 | 15.72 | **<0.001** |
| Pre-stimulus gaze direction type (NLDE) | -0.09 | 0.01 | -0.11 – -0.08 | -11.38 | **<0.001** |
| Epoch 2 | -0.01 | 0.01 | -0.03 – 0.00 | -1.50 | 0.133 |
| Epoch 3 | -0.00 | 0.01 | -0.02 – 0.01 | -0.45 | 0.651 |
| Epoch 4 | -0.00 | 0.01 | -0.02 – 0.02 | -0.14 | 0.889 |
| Group | 0.00 | 0.01 | -0.01 – 0.01 | 0.58 | 0.559 |
| Pre-stimulus gaze direction type (LDE) × Epoch 2 | -0.01 | 0.01 | -0.04 – 0.01 | -1.00 | 0.316 |
| Pre-stimulus gaze direction type (NLDC) × Epoch 2 | -0.03 | 0.01 | -0.05 – 0.00 | -1.84 | 0.066 |
| Pre-stimulus gaze direction type (NLDE) × Epoch 2 | 0.02 | 0.01 | -0.01 – 0.05 | 1.19 | 0.235 |
| Pre-stimulus gaze direction type (LDE) × Epoch 3 | -0.02 | 0.01 | -0.04 – 0.01 | -1.07 | 0.285 |
| Pre-stimulus gaze direction type (NLDC) × Epoch 3 | -0.00 | 0.01 | -0.03 – 0.03 | -0.19 | 0.846 |
| Pre-stimulus gaze direction type (NLDE) × Epoch 3 | 0.01 | 0.01 | -0.02 – 0.04 | 0.68 | 0.496 |
| Pre-stimulus gaze direction type (LDE) × Epoch 4 | 0.00 | 0.01 | -0.02 – 0.03 | 0.32 | 0.747 |
| Pre-stimulus gaze direction type (NLDC) × Epoch 4 | -0.00 | 0.01 | -0.03 – 0.03 | -0.02 | 0.986 |
| Pre-stimulus gaze direction type (NLDE) × Epoch 4 | -0.01 | 0.01 | -0.03 – 0.02 | -0.41 | 0.681 |

| | | | | | |
|---|---|---|---|---|---|
| Pre-stimulus gaze direction type (LDE) × Group | -0.00 | 0.01 | -0.02 – 0.02 | -0.03 | 0.975 |
| Pre-stimulus gaze direction type (NLDC) × Group | 0.01 | 0.01 | -0.00 – 0.03 | 1.45 | 0.148 |
| Pre-stimulus gaze direction type (NLDE) × Group | -0.01 | 0.01 | -0.02 – 0.01 | -0.70 | 0.486 |
| Epoch 2 × Group | 0.01 | 0.01 | -0.00 – 0.03 | 1.42 | 0.155 |
| Epoch 3 × Group | -0.00 | 0.01 | -0.02 – 0.01 | -0.41 | 0.684 |
| Epoch 4 × Group | -0.01 | 0.01 | -0.02 – 0.01 | -0.73 | 0.466 |
| Pre-stimulus gaze direction type (LDE) × Epoch 2 × Group | -0.01 | 0.01 | -0.04 – 0.02 | -0.70 | 0.487 |
| Pre-stimulus gaze direction type (NLDC) × Epoch 2 × Group | 0.01 | 0.01 | -0.01 – 0.04 | 0.95 | 0.344 |
| Pre-stimulus gaze direction type (NLDE) × Epoch 2 × Group | 0.02 | 0.01 | -0.01 – 0.04 | 1.13 | 0.258 |
| Pre-stimulus gaze direction type (LDE) × Epoch 3 × Group | 0.02 | 0.01 | -0.00 – 0.05 | 1.70 | 0.090 |
| Pre-stimulus gaze direction type (NLDC) × Epoch 3 × Group | -0.01 | 0.01 | -0.04 – 0.02 | -0.77 | 0.440 |
| Pre-stimulus gaze direction type (NLDE) × Epoch 3 × Group | -0.02 | 0.01 | -0.04 – 0.01 | -1.15 | 0.249 |
| Pre-stimulus gaze direction type (LDE) × Epoch 4 × Group | -0.02 | 0.01 | -0.04 – 0.01 | -1.15 | 0.250 |
| Pre-stimulus gaze direction type (NLDC) × Epoch 4 × Group | 0.01 | 0.01 | -0.02 – 0.03 | 0.39 | 0.697 |
| Pre-stimulus gaze direction type (NLDE) × Epoch 4 × Group | -0.00 | 0.01 | -0.03 – 0.03 | -0.12 | 0.908 |
| **Random Effects** | | | | | |
| $\sigma^2$ | 0.02 | | | | |
| $\tau_{00\ \text{subject number}}$ | 0.00 | | | | |
| ICC | 0.03 | | | | |
| N | 53 | | | | |

| | |
|---|---|
| Observations | 848 |
| Marginal $R^2$ / Conditional $R^2$ | 0.325 / 0.344 |

**Does theta synchronization affect a control declarative memory task?**

The ANOVA for the item score revealed a significant main effect of Time ($F_{(1, 58)} = 28.462$, $p < 0.001$, $\eta_p^2 = 0.329$). Participants, regardless of their group assignment, had a lower item learning score on the post-tACS session compared to the tACS session (M = 0.271, SE = 0.051). The Group main effect ($F_{(1, 58)} = 0.024$, $p = 0.879$, $\eta_p^2 < 0.001$), as well as the Time × Group interaction ($F_{(1, 58)} = 0.027$, $p = 0.871$, $\eta_p^2 < 0.001$) were not significant.

The ANOVA for the recollection score revealed a non-significant Time ($F_{(1, 58)} = 0.073$, $p = 0.789$, $\eta_p^2 = 0.001$), and Group main effect ($F_{(1, 58)} = 0.222$, $p = 0.639$, $\eta_p^2 = 0.004$), as well as a non-significant Time × Group interaction ($F_{(1, 58)} = 1.079$, $p = 0.303$, $\eta_p^2 = 0.018$).

#### Supplementary Tables

##### Table S1. LMM predicting trial-wise RT in the post-tACS session

| | | | **RT** | | |
|---|---|---|---|---|---|
| *Terms* | *b* | *SE b* | *95% CI* | *t* | *p* |
| (Intercept) | 332.69 | 4.34 | 324.19 – 341.19 | 76.75 | **<0.001** |
| Triplet Type | -2.99 | 1.09 | -5.12 – -0.86 | -2.75 | **0.006** |
| Epoch 6 | -2.37 | 0.86 | -4.06 – -0.69 | -2.76 | **0.006** |
| Epoch 7 | -0.76 | 0.86 | -2.45 – 0.94 | -0.88 | 0.381 |
| Group | -1.82 | 4.34 | -10.32 – 6.67 | -0.42 | 0.674 |
| Triplet Type × Epoch 6 | -0.79 | 0.86 | -2.48 – 0.90 | -0.92 | 0.359 |
| Triplet Type × Epoch 7 | 3.14 | 0.86 | 1.45 – 4.84 | 3.64 | **<0.001** |
| Triplet Type × Group | -0.99 | 1.09 | -3.12 – 1.14 | -0.91 | 0.362 |
| Epoch 6 × Group | -2.69 | 0.86 | -4.38 – -1.01 | -3.13 | **0.002** |
| Epoch 7 × Group | -0.56 | 0.86 | -2.25 – 1.14 | -0.64 | 0.519 |
| Triplet Type × Epoch 6 × Group | 1.81 | 0.86 | 0.12 – 3.49 | 2.10 | **0.036** |
| Triplet Type × Epoch 7 × Group | -1.36 | 0.86 | -3.06 – 0.33 | -1.58 | 0.114 |
| **Random Effects** | | | | | |
| $\sigma^2$ | 11923.07 | | | | |
| $\tau_{00}$ subject number | 934.25 | | | | |
| $\tau_{11}$ subject number.× Triplet type | 41.25 | | | | |
| $\rho_{01}$ subject number | 0.29 | | | | |
| ICC | 0.08 | | | | |

| | |
|---|---|
| N | 53 |
| Observations | 40924 |
| Marginal $R^2$ / Conditional $R^2$ | 0.002 / 0.079 |

**Table S2. LMM predicting trial-wise pre-stimulus gaze direction changes in the post-tACS session**

| *Terms* | *b* | *SE b* | *95% CI* | *z* | *p* |
|---|---|---|---|---|---|
| (Intercept) | -1.6308 | 0.2613 | -2.1430 – -1.1186 | -6.2407 | **<0.001** |
| Epoch 6 | 0.1347 | 0.1012 | -0.0636 – 0.3331 | 1.3312 | 0.183 |
| Epoch 7 | 0.4053 | 0.1212 | 0.1677 – 0.6428 | 3.3436 | **0.001** |
| Group | -0.8126 | 0.4172 | -1.6302 – 0.0050 | -1.9479 | 0.051 |
| Epoch 6 × Group | 0.4186 | 0.1627 | 0.0998 – 0.7374 | 2.5736 | **0.010** |
| Epoch 7 × Group | 0.3293 | 0.1958 | -0.0545 – 0.7131 | 1.6818 | 0.093 |
| **Random Effects** | | | | | |
| $\sigma^2$ | 3.29 | | | | |
| $\tau_{00\ \text{subject number}}$ | 2.14 | | | | |
| $\tau_{11\ \text{subject number.epoch6}}$ | 0.24 | | | | |
| $\tau_{11\ \text{subject number.epoch7}}$ | 0.39 | | | | |
| $\rho_{01}$ | -0.50 | | | | |
| | -0.67 | | | | |
| ICC | 0.34 | | | | |
| N | 53 | | | | |
| Observations | 40924 | | | | |
| Marginal $R^2$ / Conditional $R^2$ | 0.026 / 0.358 | | | | |

**Table S3. LMM predicting trial-wise learning-dependent anticipations in the post-tACS session**

| *Terms* | *b* | *SE b* | *95% CI* | *z* | *p* |
|---|---|---|---|---|---|
| (Intercept) | -2.7537 | 0.2265 | -3.1977 – -2.3096 | -12.1551 | **<0.001** |
| Epoch 6 | -0.2014 | 0.0987 | -0.3948 – -0.0081 | -2.0416 | **0.041** |
| Epoch 7 | 0.4267 | 0.1055 | 0.2199 – 0.6336 | 4.0437 | **<0.001** |
| Group | -0.5903 | 0.3596 | -1.2950 – 0.1145 | -1.6417 | 0.101 |
| Epoch 6 × Group | 0.2883 | 0.1481 | -0.0020 – 0.5785 | 1.9466 | 0.052 |
| Epoch 7 × Group | 0.1161 | 0.1669 | -0.2111 – 0.4434 | 0.6957 | 0.487 |
| **Random Effects** | | | | | |
| $\sigma^2$ | 3.29 | | | | |
| $\tau_{00\ \text{subject number}}$ | 1.52 | | | | |
| $\tau_{11\ \text{subject number.epoch6}}$ | 0.11 | | | | |
| $\tau_{11\ \text{subject number.epoch7}}$ | 0.19 | | | | |
| $\rho_{01}$ | -0.61 | | | | |
| | -0.74 | | | | |
| ICC | 0.27 | | | | |
| N | 53 | | | | |
| Observations | 40924 | | | | |
| Marginal $R^2$ / Conditional $R^2$ | 0.025 / 0.284 | | | | |

**Table S4. LMM predicting epoch-wise learning-dependent anticipation ratio in the post-tACS session**

| *Terms* | *b* | *SE b* | *95% CI* | *t* | *p* |
|---|---|---|---|---|---|
| (Intercept) | 38.30 | 1.25 | 35.84 – 40.77 | 30.58 | **<0.001** |
| Epoch 6 | 4.16 | 1.79 | 0.64 – 7.67 | 2.32 | **0.021** |
| Epoch 7 | -5.34 | 1.53 | -8.34 – -2.34 | -3.50 | **0.001** |
| Group | 0.78 | 1.25 | -1.69 – 3.24 | 0.62 | 0.536 |
| Epoch 6 × Group | -0.87 | 1.79 | -4.38 – 2.65 | -0.48 | 0.628 |
| Epoch 7 × Group | -0.71 | 1.53 | -3.71 – 2.29 | -0.47 | 0.642 |
| **Random Effects** | | | | | |
| $\sigma^2$ | 504.38 | | | | |
| $\tau_{00\ \text{subject number}}$ | 6.45 | | | | |
| ICC | 0.01 | | | | |
| N | 53 | | | | |
| Observations | 402 | | | | |
| Marginal $R^2$ / Conditional $R^2$ | 0.036 / 0.048 | | | | |

**Table S5. LMM predicting trial-wise reaction time as a function of information type (LL, HL, LH, HH) on epoch 6**

| *Terms* | *b* | *SE b* | *95% CI* | *t* | *p* |
|---|---|---|---|---|---|
| (Intercept) | 286.49 | 3.43 | 279.76 – 293.21 | 83.51 | **<0.001** |

| | | | | | |
|---|---|---|---|---|---|
| Information type (HL) | 11.40 | 2.12 | 7.26 – 15.55 | 5.39 | **<0.001** |
| Information type (LH) | 8.06 | 3.07 | 2.06 – 14.07 | 2.63 | **0.009** |
| Information type (HH) | -6.72 | 1.86 | -10.36 – -3.08 | -3.61 | **<0.001** |
| Group | 1.45 | 3.43 | -5.28 – 8.17 | 0.42 | 0.673 |
| Information type (HL) × Group | 2.41 | 2.12 | -1.74 – 6.56 | 1.14 | 0.255 |
| Information type (LH) × Group | 0.58 | 3.07 | -5.43 – 6.59 | 0.19 | 0.851 |
| Information type (HH) × Group | 3.34 | 1.86 | -0.30 – 6.99 | 1.80 | 0.072 |
| **Random Effects** | | | | | |
| $\sigma^2$ | 27816.53 | | | | |
| $\tau_{00\ \text{subject number}}$ | 496.12 | | | | |
| ICC | 0.02 | | | | |
| $N_{\text{subject number}}$ | 53 | | | | |
| Observations | 20962 | | | | |
| Marginal $R^2$ / Conditional $R^2$ | 0.004 / 0.021 | | | | |

**Table S6. LMM predicting trial-wise learning-dependent anticipation as a function of information type (LL, HL, LH, HH) on epoch 6**

| *Terms* | *b* | *SE b* | *95% CI* | *t* | *p* |
|---|---|---|---|---|---|
| (Intercept) | -2.99 | 0.22 | -3.42 – -2.57 | -13.81 | **<0.001** |
| Information type (HL) | -0.27 | 0.14 | -0.55 – 0.00 | -1.96 | 0.050 |
| Information type (LH) | -0.18 | 0.08 | -0.34 – -0.01 | -2.13 | **0.033** |
| Information type (HH) | 0.53 | 0.10 | 0.33 – 0.73 | 5.29 | **<0.001** |
| Group | -0.34 | 0.35 | -1.02 – 0.34 | -0.97 | 0.331 |
| Information type (HL) × Group | 0.29 | 0.23 | -0.17 – 0.75 | 1.22 | 0.221 |

| | | | | | |
|---|---|---|---|---|---|
| Information type (LH) × Group | 0.34 | 0.14 | 0.06 – 0.62 | 2.37 | **0.018** |
| Information type (HH) × Group | -0.72 | 0.19 | -1.10 – -0.34 | -3.73 | **<0.001** |
| **Random Effects** | | | | | |
| $\sigma^2$ | 3.29 | | | | |
| $\tau_{00\ \text{subject number}}$ | 1.32 | | | | |
| ICC | 0.29 | | | | |
| $N_{\text{subject number}}$ | 53 | | | | |
| Observations | 20962 | | | | |
| Marginal $R^2$ / Conditional $R^2$ | 0.013 / 0.295 | | | | |

**Table S7. LMM predicting epoch-wise likelihood of pre-stimulus gaze direction types as a function of pre-stimulus gaze direction types (LDC, LDE, NLDC, NLDE), epoch, and group**

| *Terms* | *b* | *SE b* | *95% CI* | *t* | *p* |
|---|---|---|---|---|---|
| (Intercept) | 1.04 | 0.01 | 1.03 – 1.06 | 171.31 | **<0.001** |
| LDE | 0.04 | 0.01 | 0.02 – 0.06 | 3.97 | **<0.001** |
| NLDC | 0.13 | 0.01 | 0.11 – 0.15 | 13.53 | **<0.001** |
| NLDE | -0.08 | 0.01 | -0.10 – -0.06 | -8.85 | **<0.001** |
| Epoch 6 | 0.01 | 0.01 | -0.01 – 0.02 | 0.99 | 0.321 |
| Epoch 7 | -0.00 | 0.01 | -0.02 – 0.01 | -0.28 | 0.776 |
| Group | 0.01 | 0.01 | 0.00 – 0.03 | 2.29 | **0.022** |
| LDE × Epoch 6 | 0.01 | 0.01 | -0.02 – 0.03 | 0.42 | 0.671 |
| NLDC × Epoch 6 | -0.02 | 0.01 | -0.04 – 0.01 | -1.40 | 0.162 |
| NLDE × Epoch 6 | 0.03 | 0.01 | 0.00 – 0.05 | 2.04 | **0.042** |
| LDE × Epoch 7 | -0.01 | 0.01 | -0.04 – 0.02 | -0.77 | 0.444 |
| NLDC × Epoch 7 | 0.02 | 0.01 | -0.01 – 0.05 | 1.49 | 0.138 |

| | | | | | |
|---|---|---|---|---|---|
| NLDE × Epoch 7 | -0.01 | 0.01 | -0.04 – 0.01 | -1.10 | 0.273 |
| LDE × Group | 0.01 | 0.01 | -0.01 – 0.03 | 0.71 | 0.476 |
| NLDC × Group | 0.03 | 0.01 | 0.02 – 0.05 | 3.64 | **<0.001** |
| NLDE × Group | -0.02 | 0.01 | -0.03 – 0.00 | -1.64 | 0.101 |
| Epoch 6 × Group | -0.01 | 0.01 | -0.02 – 0.01 | -1.27 | 0.205 |
| Epoch 7 × Group | 0.00 | 0.01 | -0.01 – 0.02 | 0.16 | 0.873 |
| LDE × Epoch 6 × Group | -0.01 | 0.01 | -0.04 – 0.01 | -0.92 | 0.360 |
| NLDC × Epoch 6 × Group | 0.01 | 0.01 | -0.02 – 0.03 | 0.41 | 0.679 |
| NLDE × Epoch 6 × Group | -0.01 | 0.01 | -0.04 – 0.01 | -0.90 | 0.366 |
| LDE × Epoch 7 × Group | 0.01 | 0.01 | -0.02 – 0.04 | 0.79 | 0.431 |
| NLDC × Epoch 7 × Group | -0.02 | 0.01 | -0.04 – 0.01 | -1.31 | 0.191 |
| NLDE × Epoch 7 × Group | 0.01 | 0.01 | -0.02 – 0.04 | 0.77 | 0.440 |
| **Random Effects** | | | | | |
| $\sigma^2$ | 0.02 | | | | |
| $\tau_{00}$ subject number | 0.00 | | | | |
| ICC | 0.02 | | | | |
| N subject number | 53 | | | | |
| Observations | 636 | | | | |
| Marginal $R^2$ / Conditional $R^2$ | 0.346 / 0.359 | | | | |
